# cisDynet: an integrated platform for modeling gene-regulatory dynamics and networks

**DOI:** 10.1101/2023.10.30.564662

**Authors:** Tao Zhu, Xinkai Zhou, Yuxin You, Lin Wang, Zhaohui He, Dijun Chen

**Author notes:** Correspondence (Dijun Chen). These authors contributed equally to this work (Tao Zhu, Xinkai Zhou, Yuxin You).

## Abstract

Chromatin accessibility sequencing has been widely used for uncovering genetic regulatory mechanisms and inferring gene regulatory networks. However, effectively integrating large-scale chromatin accessibility datasets has posed a significant challenge. This is due to the lack of a comprehensive end-to-end solution, as many existing tools primarily emphasize data pre-processing and overlook downstream analyses. To bridge this gap, we have introduced cisDynet, a holistic solution that combines streamlined data pre-processing using Snakemake and R functions with advanced downstream analysis capabilities. cisDynet excels in conventional data analyses, encompassing peak statistics, peak annotation, differential analysis, motif enrichment analysis and more. Additionally, it allows to perform sophisticated data exploration such as tissue-specific peak identification, time-course data fitting, integration of RNA-seq data to establish peak-to-gene associations, constructing regulatory networks, and conducting enrichment analysis of GWAS variants. As a proof of concept, we applied cisDynet to re-analyze the comprehensive ATAC-seq datasets across various tissues from the ENCODE project. The analysis successfully delineated tissue-specific open chromatin regions (OCRs), established connections between OCRs and target genes, and effectively linked these discoveries with 1,861 GWAS variants. Furthermore, cisDynet was instrumental in dissecting the time-course open chromatin data of mouse embryonic development, revealing the dynamic behavior of OCRs over time and identifying key transcription factors governing differentiation trajectories. In summary, cisDynet offers researchers a user-friendly solution that minimizes the need for extensive coding, ensures the reproducibility of results, and greatly simplifies the exploration of epigenomic data.

**Graphical Abstract:** cisDynet enables the exploration of the cis-regulatory chromatin dynamics and networks. It offers a streamlined preprocessing pipeline built on Snakemake, along with an R package for advanced downstream data analysis and visualization. It is free to access on GitHub (https://github.com/tzhu-bio/cisDynet).

**Highlights:**

- The cisDynet enables comprehensive and efficient processing of chromatin accessibility data, including pre-processing, advanced downstream data analysis and visualization.
- cisDynet provides a range of analytical features such as processing of time course data, co-accessibility analysis, linking OCRs to genes, building regulatory networks, and GWAS variant enrichment analysis.
- cisDynet simplifies the identification of tissue/cell type-specific OCRs or dynamic OCR changes over time and facilitates the integration of RNA-seq data to depict temporal trajectories.

## INTRODUCTION

In the nuclei of eukaryotic cells, DNA is intricately packaged and surrounded by nucleosomes. Regions of the genome that lack nucleosomes are known as chromatin accessible or open chromatin regions (OCRs). These OCRs serve as binding sites for transcription factors (TF) and RNA polymerases, playing a crucial role in regulating downstream gene expression [1, 2]. To identify OCRs within the genome, various high-throughput genomics technologies are available, including ATAC-seq [3, 4], MNase-seq [5], FAIRE-seq [6], and DNase-seq [7, 8]. Among these, ATAC-seq has gained widespread popularity due to its advantages, such as shorter experimental timeline, minimal requirements for starting material, and enhanced sensitivity [4, 9].

Standard analysis procedures for ATAC-seq and other chromatin accessibility data typically involve tasks such as peak identification and annotation, differential peak analysis, footprint analysis, and the construction of regulatory networks [9]. To carry out these analyses efficiently, researchers often need to employ multiple software packages, each with its specific input requirements. This can be somewhat inconvenient and challenging. As of now, there is a noticeable gap in the availability of a comprehensive toolkit for analyzing chromatin accessibility data that can seamlessly accommodate a wide range of usage scenarios.

The assignment of target genes is a pivotal step in the analysis of chromatin accessibility data. In cases where datasets have limited sample sizes, gene annotation is typically performed by associating the summits of OCRs with the nearest transcription start site (TSS). Once the target genes for differential peaks are identified, they are subsequently compared with the differentially expressed genes obtained from RNA-seq data for joint analysis. Several software packages, including DiffBind [10], HOMER [11] and others, already support this type of analysis. In the context of analyzing multiple samples, assigning target genes can be accomplished by considering the correlation of chromatin accessibility and gene expression across different samples. This approach has been proven to be effective and feasible [12, 13]. However, it is worth noting that there is currently no dedicated tool available specifically designed for conducting this type of analysis.

Here we created cisDynet, a user-friendly and all-encompassing toolkit designed for modeling gene-regulatory dynamics and networks using chromatin accessibility data. This toolkit consists of two essential components: a data pre-processing workflow based on Snakemake [14] and an R package that provides a wide range of functions and visualization tools. In addition to standard analyses like peak statistical assessment, differential peak analysis, motif and footprint evaluation, we have incorporated advanced features tailored to handle multiple samples effectively. These features encompass the analysis of time-course data, co-accessibility analysis, linking OCRs to specific genes, constructing regulatory networks, performing GWAS variant enrichment analyses, and more. Importantly, these functions demand minimal computational resources and are compatible with personal computers. To facilitate user understanding and implementation of these functions, we have provided a comprehensive manual accessible at (https://tzhu-bio.github.io/cisDynet_bookdown/book/index.html). Furthermore, we have applied cisDynet to reanalyze the ATAC-seq data for 17 human tissues obtained from ENCODE project. Our analysis successfully identified tissue-specific OCRs, established connections between OCRs and target genes, and effectively associated these findings with 1,861 GWAS variants. Additionally, we employed cisDynet to analyze time-course data from mouse embryo development, revealing dynamic changes in OCRs over time and identifying key transcription factors governing the trajectory of differentiation. In summary, cisDynet streamlines the process of chromatin accessibility data analysis, reducing the need for extensive code development, and ensuring the reproducibility of results.

## RESULTS

### Overview of cisDynet

cisDynet comprises two core components: a data preprocessing workflow, implemented based on Snakemake [14], and an accompanying R package designed for subsequent downstream analysis. Our data preprocessing workflow adheres to the established standard reference process outlined by the ENCODE project [15]. This comprehensive process encompasses tasks such as sequencing adapter removal, read alignment, data filtering, quality control (QC) metrics calculation, and peak calling, among others. A notable feature of our data preprocessing process is its adaptability in effectively eliminating true PCR duplicates through the utilization of UMI-based sequencing [16]. Upon completion of this process, cisDynet automatically generates a comprehensive HTML report. This report offers users a clear and informative overview of the data quality assessment results, including essential metrics such as Transcription Start Site (TSS) enrichment scores and FRiP (Fraction of Reads in Peaks) (Supplementary Fig. 1). These metrics are widely recognized benchmarks for evaluating data quality in epigenomics. Additionally, cisDynet provide insertion fragment distributions specifically tailored for chromatin accessibility data. To ensure high data quality, the insert fragment distribution should exhibit a distinct nucleosome pattern and a notable enrichment in proximity to the TSS (Supplementary Fig. 2).

The cisDynet R package offers a comprehensive suite of user-friendly functions that cover a broad spectrum of tasks, ranging from quality control to advanced downstream analysis and visualization (Fig. 1). Detailed tutorials are available at https://tzhu-bio.github.io/cisDynet_bookdown/book/index.html. From a global perspective, the cisDynet package empowers users to effortlessly perform a diverse array of analyses, including: (1) assessing similarity between different sets of peaks; (2) annotating peaks; (3) conducting two-groups/three-groups differential peak analysis; (4) performing differential analysis across multiple samples; (5) fitting time-course data and integrating RNA-seq data for the identification of dynamically changing OCRs and associated genes; (6) quantifying TF activity through footprinting analysis; (7) conducting co-accessibility analysis of OCRs; (8) exploring genome-wide associations between peaks and target genes; (9) constructing TF-TF regulatory networks; (10) linking GWAS (Genome-Wide Association Study) traits through enrichment analysis of genetic variations. Furthermore, our package offers methods for tasks such as super-enhancer identification and enrichment analysis among sets of peaks. In summary, cisDynet provides a wealth of user-friendly functions that facilitate a wide range of analyses, making it particularly valuable for researchers focused on unraveling the intricacies of gene expression regulation through open chromatin regions and related investigations.

**Figure 1.**
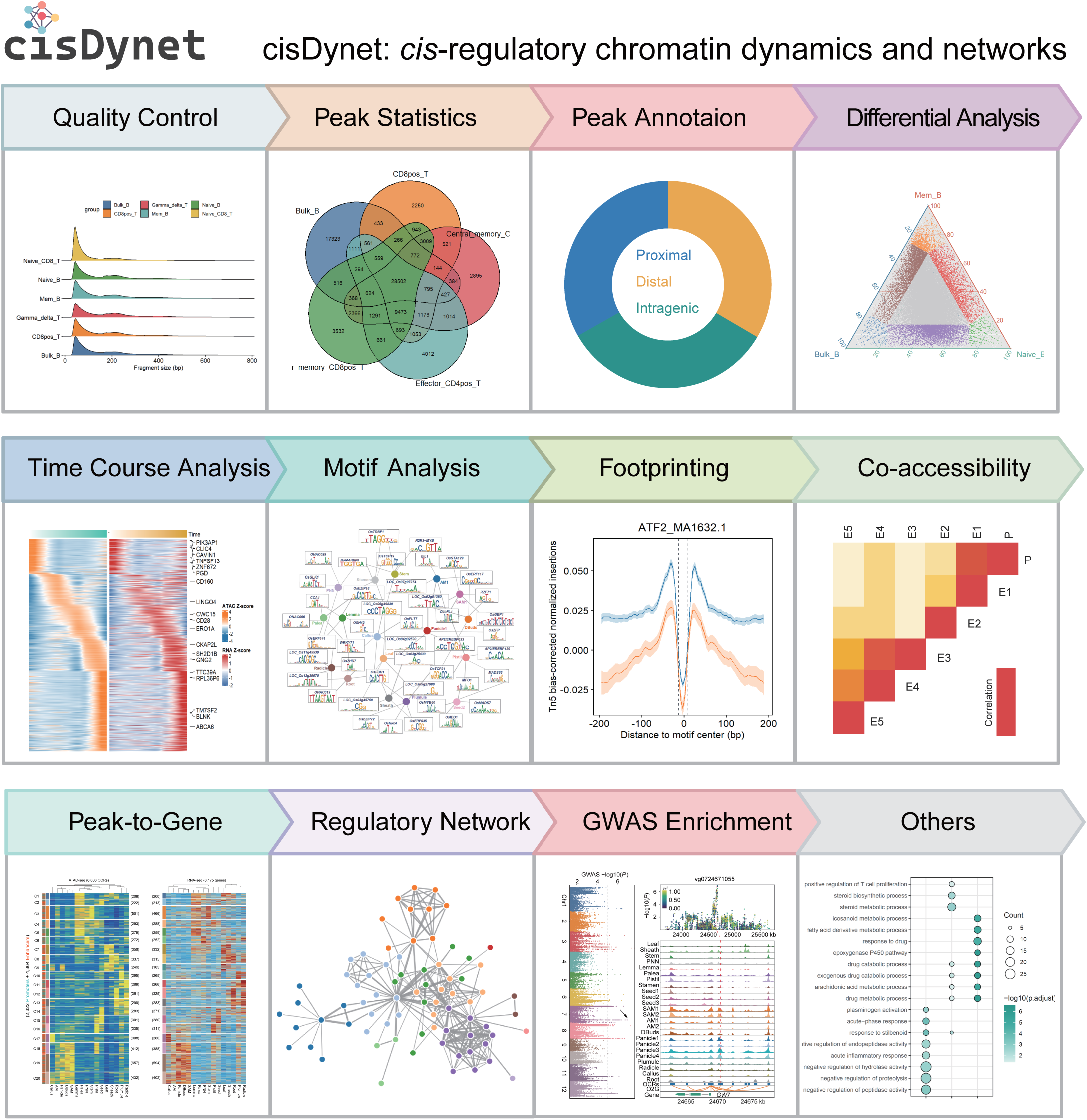
Overview of the functional modules for analyzing chromatin accessibility data provided by the cisDynet R package.

### Comparison of cisDynet with other tools

In the arena of ATAC-seq data processing, several software tools have developed, including esATAC [17], ATACseqQC [18], AIAP [19], and PEPATAC [20]. These tools primarily serve the purpose of preprocessing ATAC-seq data, which involves tasks like removing sequencing adapters, aligning reads to the genome, and calling peaks. However, their primary concentration remains confined to these foundational data processing phases (Fig. 2). They tend to have limited capabilities for carrying out downstream advanced analyses, such as conducting differential analysis of multiple samples, linking regulatory regions to gene expression, constructing regulatory networks, exploring the enrichment of GWAS variants, and so on (Fig. 2b).

**Figure 2.**
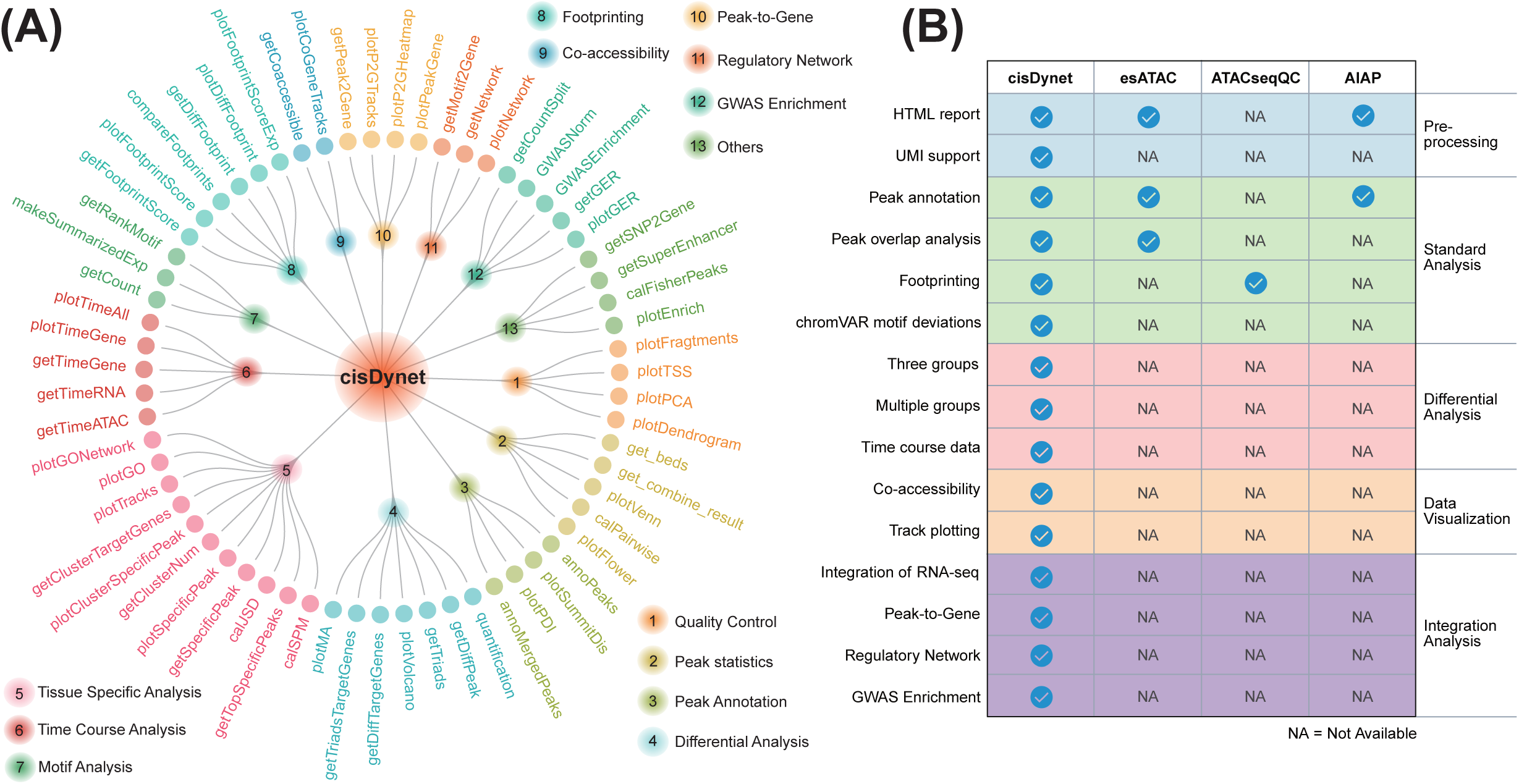
(A) The functions for the different modules provided by cisDynet. (B) Comparison of the cisDynet with other tools used to process chromatin accessibility data.

We have developed cisDynet to provide an efficient analytical pipeline that covers the entire spectrum of ATAC-seq data analyses, from essential data pre-processing to comprehensive downstream advanced analysis (Fig. 2a, Supplementary Fig. 3). cisDynet is meticulously designed to accommodate a wide array of ATAC-seq analysis scenarios, effectively catering to the diverse needs of researchers. A significant feature of cisDynet is its capability to seamlessly integrate data from ATAC-seq and RNA-seq for joint analysis -- a functionality that is seldom addressed in existing software. Traditionally, most ATAC-seq tools follow an independent process: they firstly identify differential OCRs and differential genes separately and then attempt to connect OCRs to the nearest genes before performing an intersection with differential genes. However, this approach becomes less effective when dealing with large sample sizes or population-scale data. In this context, our cisDynet analysis pipeline offers a unique advantage. It not only identifies differential OCRs and genes but also seamlessly integrates both ATAC-seq and RNA-seq data. Furthermore, while TOBIAS [21] currently represents a featured tool for construction of regulatory network based ATAC-seq data, it focuses on utilizing footprint to construct TF-TF regulatory network without integrating RNA-seq data. In contrast, cisDynet builds on this foundation by integration of RNA-seq data, thereby enhancing the confidence of the regulatory network construction (Supplementary Fig. 4).

The presence of some anomalous regions on the genome allows for extremely high signal. Most of these regions are due to the presence of repetitive sequences, genome assembly errors, etc. In mammals, “problematic” regions have been inferred and manually checked and called blacklists, and are widely used in the analysis of genomic data. However, there is no systematic blacklists for plant genomes at present. To fill this gap, we used the greenscreen software[22] in combination with data collected from our ChIP-Hub database [23] to obtain “problematic” lists for five plant species: Arabidopsis thaliana, rice, maize, soybean, and tomato. We succeeded in identifying “problematic” regions in Arabidopsis thaliana and rice that may be due to genome assembly errors (Supplementary Fig. 5). We believe that excluding these blacklisted regions when analyzing plant epigenomic data using cisDynet will improve quantitative accuracy.

### cisDynet provides comprehensive differential analyses of multiple samples

The identification of differential peaks serves a dual purpose: it aids in discerning the impact of specific conditions or treatments on the chromatin states of genes and also facilitates the identification of potential regulators associated to these differential peaks. In addition to the widely-used two-cohort differential peaks identification provided by cisDynet, our package offers advanced three-cohort and multi-cohort differential peak analysis with accompanying visualization capabilities (Fig. 3). For the analysis of differential peaks across the three cohorts, we drew inspiration from a previous study [24] and devised a triad analysis strategy. Initially, we construct a pseudo-matrix comprising seven subgroups: A dominant, B dominant, C dominant, A suppressed, B suppressed, C suppressed, and balanced (Fig. 3A). Subsequently, for the quantitatively normalized matrix, we calculated the Euclidean distance between the quantitative results of each peak and each of the seven subgroupings within that pseudo-matrix. We then select the grouping with the closest Euclidean distance to categorize that particular peak. To visualize the obtained grouping result, we employ a ternary graph. To illustrate this approach, we obtained human chromatin accessibility data from ENCODE [15], focusing on the lungs, liver, and spleen to assess differences in chromatin accessibility across these three organs (Fig. 3B). Remarkably, the differential groupings we derived align precisely with the observed quantitative distinctions, thereby validating the accuracy of our categorizations (Fig. 3C).

**Figure 3.**
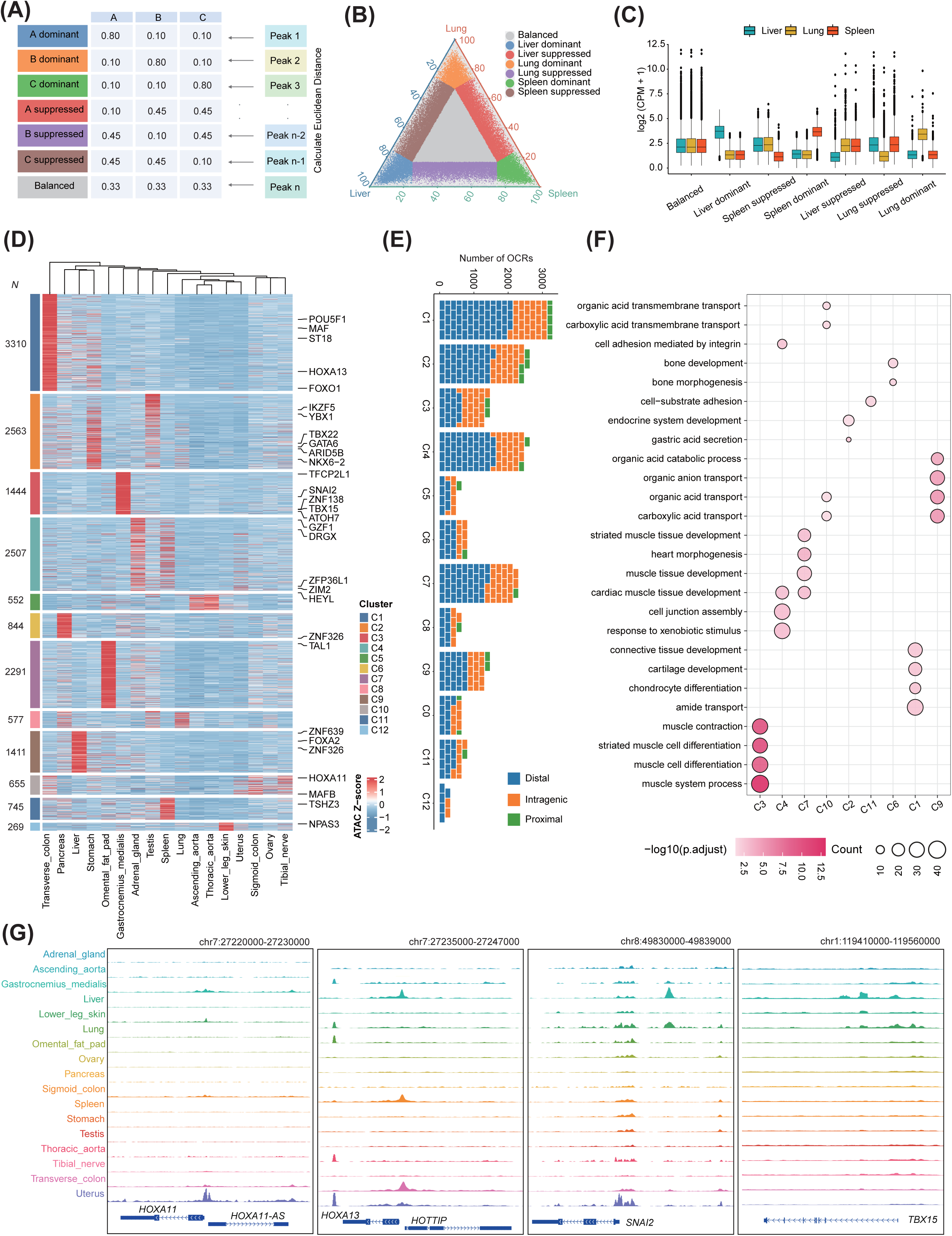
Example of using cisDynet to compare chromatin accessibility differences and identify tissue-specific OCRs. (A) Seven types of OCR predefined matrices were used to categorize chromatin accessibility differences among the three samples. (B) Ternary diagram showing chromatin accessibility differences in three organs, Lung, Liver and Spleen. (C) The boxplot showing the degree of chromatin accessibility for the seven classifications in (B). (D) The heatmap showing the tissue-specific OCRs identified using cisDynet All tissue-specific OCRs were grouped into 12 clusters according to their similarity in chromatin accessibility. (E) The proportion of intragenic, proximal, and distal OCRs included in each cluster in the (D). (F) The dotplot demonstrates the enrichment analysis of the biological pathways involved in the target genes of the OCR for each cluster. (G) The genome browser demonstrates the distribution of ATAC-seq signaling near *HOXA11*, *HOXA13*, *SNAI2* and *TBX15* loci.

For the differential analysis among more than three groups, an index of Shannon entropy is calculated for each OCR from the normalized matrix. This entropy index is used to filter and identify peaks that are highly variable among different groups. It is worth noting that this screening method may result in an unbalanced number of differential peaks among samples especially when the majority of the samples show a relatively high degree of similarity while a few samples are significantly different from the others. To illustrate this challenge, we utilized ATAC-seq data from multiple immune cell lines [25]. Significant differentiation was observed in plasmacytoid dendritic cells (pDCs) and monocytes compared to other cell lines (Supplementary Fig. 6A). Since the majority of samples consisted of T cells, our analysis mainly yielded differential peaks associated with pDCs and monocytes, making it difficult to discern variations among distinct T cell populations (Supplementary Fig. 6B). To address this challenge, we incorporated an additional metric called the specificity measure (SPM). SPM allows to discern and filter highly accessible OCRs in individual samples or across multiple samples (Supplementary Fig. 6C-6D). As expected, this approach significantly improves our ability to compare accessibility differences among samples (Supplementary Fig. 6E). We applied this strategy to the ENCODE data [15] to identify differential OCRs between distinct organs (Fig. 3D). These organ-specific OCRs predominantly located in TSS-distal regions, which is similar to previous study [23, 26] (Fig. 3E). CisDynet also provides functionality to investigate the biological pathways associated with these differential OCRs (Fig. 3D, F). The signal files provided by cisDynet can be directly visualized by epigenome browsers such as WashU Epigenome Browser [27]. For instance, Homeobox protein Hox-A13 (*HOXA11*) has been previously documented as a gene specifically expressed in epithelial and stromal cell types within the uterus [28] (Fig. 3G). Homeobox protein Hox-A13 (*HOXA13*) is another gene with highly specific expression in the colon, affecting epidermal differentiation [29] (Fig. 3G). Additional examples include Zinc finger protein SNAI2 (*SNAI2*) and T-box transcription factor TBX15 (*TBX15*), both exhibiting specific openness in the gastrocnemius medialis (Fig. 3G).

### Depicting time trajectory data with cisDynet

Time-course experiments are essential for uncovering the dynamic aspects of cellular processes, such as cell differentiation or responses to stress. Unlike single-cell pseudo-time analyses, bulk data do not require the inference of their temporal order. We focus solely on fitting dynamically changing OCRs in accordance with the time points defined in the experimental design. Firstly, the highly dynamic OCRs across different time points can be identified using the differential analysis method mentioned above. Then, we apply local polynomial regression fitting to establish the relationship between time points and the activity of these OCRs. This fitting allows us to predict the smooth change in OCR values over time based on the fitted model (Fig. 4A). As a proof of concept, we conducted an analysis using ATAC-seq data collected from five time points during the differentiation of mouse embryonic stem cells (mESCs) [30] (Fig. 4B). In total, we obtained 2,165 OCRs that changed dynamically over time and integrated RNA-seq data to find the corresponding target genes of these OCRs (Fig. 4C). Through this analysis, we observed the involvement of genes such as SRY (sex determining region Y)-box 2 (*Sox2*) [31] and Atypical chemokine receptor 3 (*Ackr3*) [32], which are known to play roles in maintaining the activity of neural stem and progenitor cells (Fig. 4D). Conversely, genes like SRY-Box Transcription Factor 11 (*Sox11*) and Glial fibrillary acidic protein (*Gfap*) play significant roles in the mid- and late-stages, respectively, with *Gfap* notably serving as a marker gene for astrocytes within the central nervous system [33] (Fig. 4D).

**Figure 4.**
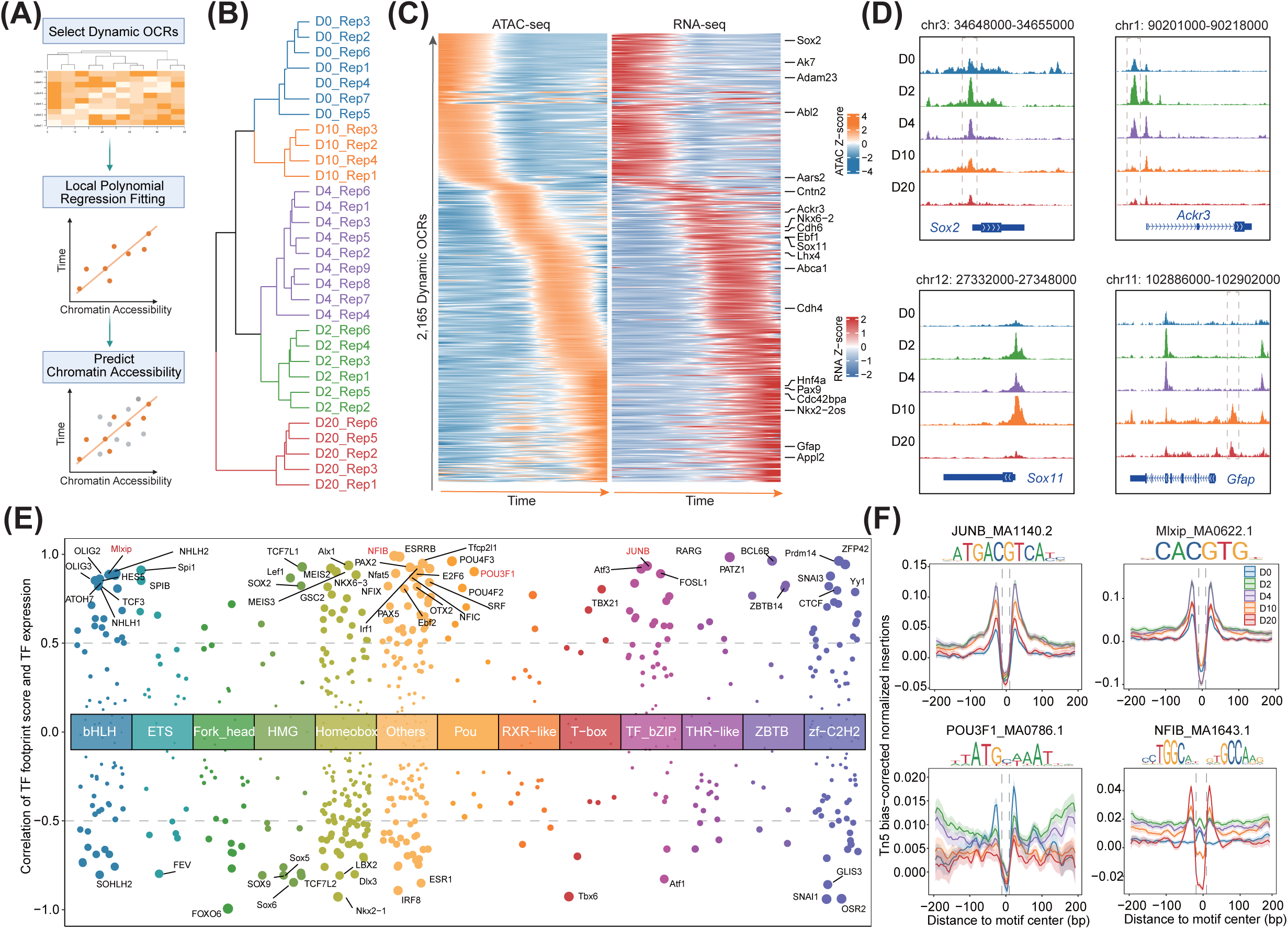
Integrating RNA-seq to identify OCRs that change dynamically over time and important TFs involved in mESCs developmental process by cisDynet. (A) Flowchart of cisDynet identification of dynamically changing OCR over time. (B) Clustering results for chromatin accessibility data at different time points. (C) The 2,165 OCRs identified using cisDynet that dynamically change over time, and the expression changes of their target genes. The target genes were identified based on the principle that they were closest to the OCRs and had a Pearson correlation coefficient of not less than 0.7 with their expression. (D) Genome browser track plot demonstrating the dynamics of OCR near *Sox2*, *Ackr3*, *Sox11* and *Gfap* genes during mESCs differentiation. (E) The scatterplot showing the distribution of Pearson correlation coefficients between the footprint score of TFs from different families and their own expression. Families with less than 15 TFs were combined into others. The TF footprint score was calculated by TOBIAS[21]. (F) Distribution of Tn5 cuts at TF footprint sites at four time points during the differentiation of mESCs. Specific TF information and motif logos are labeled above each subfigure.

mESCs differentiate into three subpopulations after two days, prompting us to investigate the key regulatory factors driving this process [30]. We first utilized TOBIAS [21] to measure the genome-wide activity of all TFs in these five stages and termed it as the footprint score. We then calculated the Pearson correlation coefficient between the footprint score and the gene expression and identified driver regulators based on the correlation coefficient (Fig. 4E). We compared TF footprint activity across five time points and observed that JUNB and MLX Interacting Protein (Mlxip) had higher activity at day 2 (D2) and D4. As we expected, it was also observed that JUNB and Mlxip had deeper footprints in the D2 and D4, respectively (Fig. 4F). Additionally, we observed strong activity of POU Class 3 Homeobox 1 (POU3F1) [34] during the early stage (D0), a gene known to be associated with neural development. Conversely, Nuclear Factor I B (NFIB) displayed remarkably high activity at D20 and was also identified as a driver regulator in the original study [30].

### Regulatory regions affecting gene expression can be effectively identified by cisDynet

The chromatin accessibility of OCRs directly impacts the expression levels of genes, and extensive studies have consistently demonstrated a strong positive correlation between the two [1, 9, 35]. This prompted us to predict potential target genes for OCR at a genome-wide scale. Due to the spatial conformation of chromatin, OCRs do not linearly regulate nearest genes. Based on information from 3D interactions, we can limit the maximum regulatory distance from the TSS (e.g., 500kb upstream and downstream in humans) (Fig. 5A). Our cisDynet can efficiently help to find potential OCRs for each gene within a specified range. Taking the ENCODE data as an example, we confined the potential regulatory region to 500 kb. CisDynet can efficiently computed peak-to-gene links in just 40 minutes with a single thread and minimal memory consumption. We identified a total of 236,792 reliable peak-to-gene links with absolute Pearson correlation coefficient (|*R*|) > 0.4 and FDR < 0.05 (Fig. 5B). As we expected, the correlation coefficient decreases with the increasing distance between OCRs and the TSS (Supplementary Fig. 7A). Furthermore, our findings indicate that peak-to-gene links are more commonly derived from proximal OCRs. On average, each gene is associated with 7.96 OCRs, whereas each OCR is associated with an average of 2.78 genes (Supplementary Fig. 7B-D). Interestingly, we noticed that TFs exhibit a higher number of peak-to-gene links compared to genes, implying that TFs may undergo more intricate regulation (Supplementary Fig. 7E). Meanwhile, for those genes or TFs with more peak-to-gene links, they are mainly enriched for “differentiation” and “development” biological pathways (Supplementary Fig. 7F).

**Figure 5.**
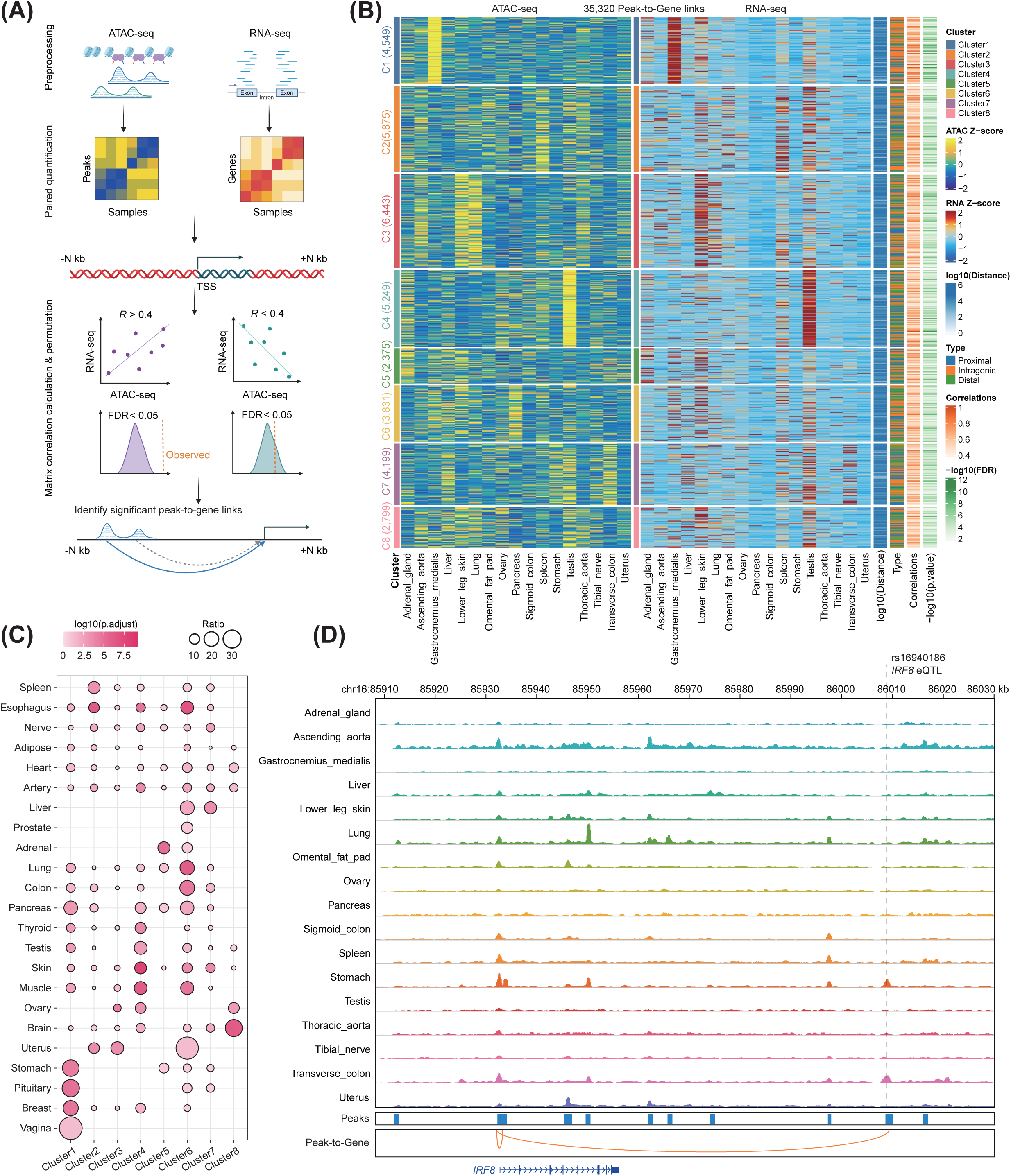
Assigning OCRs to target genes. (A) Flowchart of the computational method for establishing the correlation between OCRs and target genes. (B) The heatmap shows 35,230 peak-to-gene links computed by cisDynet (*R* > 0.4, FDR < 0.05). Only peak-to-gene links within 50 kb upstream and downstream of each gene are shown. Each row represents an OCR or gene. All OCRs are grouped into 8 clusters based on the similarity of chromatin accessibility. (C) The dotplot showing the enrichment of peak-to-gene links in each cluster in (B) with published eQTL data. The degree of enrichment was calculated by a hypergeometric test. (D) The genome browser screenshot demonstrates the presence of an eQTL (rs16940186) about 80kb downstream of *IRF8*, which happens to fall in the OCR where the peak-to-gene is located.

To further validate the reliability of our peak-to-gene linkages, we co-localized the obtained peak-to-gene links with eQTL data from various human tissues [36]. We found these identified peak-to-gene links were significantly enriched for eQTL and a tissue-specific enrichment pattern could be observed (Fig. 5C). As an example, there is an eQTL (rs16940186) located approximately 80 kb downstream of *IRF8*. Previous studies have demonstrated that this eQTL plays an important role mainly in the stomach epithelial cell type [37]. We identified peak-to-gene links in which Interferon regulatory factor 8 (*IRF8*) coincidentally associates with the OCR where this eQTL is located (*R* = 0.61, FDR = 3.8e-03), and the OCR was specific accessible in the stomach (Fig. 5D).

### cisDynet streamlines the analysis of non-coding GWAS variants

Genome-wide association studies (GWAS) have been instrumental in identifying numerous loci or genes associated with various phenotypes and diseases. However, a substantial portion of these associated loci is often located in non-coding regions of the genome [38–40]. Currently, there are several tools and resources available to associate tissue/cell types with diseases and traits by assessing whether causal variants identified in GWAS are enriched in regulatory elements [41]. These advances help bridge the gap between genetic variants and their functional impact in specific biological contexts. However, these enrichment tools predominantly rely on the mere overlap between peaks and variants, lacking consideration for quantitative information regarding the OCRs and demonstrating limited sensitivity in capturing subtle sample differences. To address this issue, Soskic et al. developed a tool named CHEERS [42], which associates traits with cell types based on quantitative information from peaks. This approach aligns with earlier findings indicating that GWAS-related variants often fall in tissue/cell type-specific regulatory regions [38, 43]. Following this principle, we introduced tissue-specific scores in the cisDynet to accommodate analysis scenarios characterized by subtle difference among samples (Fig. 6A). To validate the power of the method, we obtained data from the NHGRI-EBI GWAS Catalog [44] and filtered the GWAS with the number of significant variants associated to no less than 20 and finally obtained 2,498 GWAS. We used ENCODE chromatin accessibility data for different tissues to establish tissue associations with different GWAS variants. Throughout these enrichment results, we found that 1,861 (74.5%) GWAS were significantly enriched in at least one tissue (*P* < 0.05) and indeed showed a tissue-specific pattern (Fig. 6B). And in the absence of any prior knowledge, such associations are to some extent in accordance with current biological knowledge. For instance, COVID-19 is associated with the lung, fatty liver is associated with the liver, and ovarian cancer is associated with the ovaries (Fig. 6C). These results demonstrate the ability of cisDynet to establish associations between tissue/cell types and GWAS variants.

**Figure 6.**
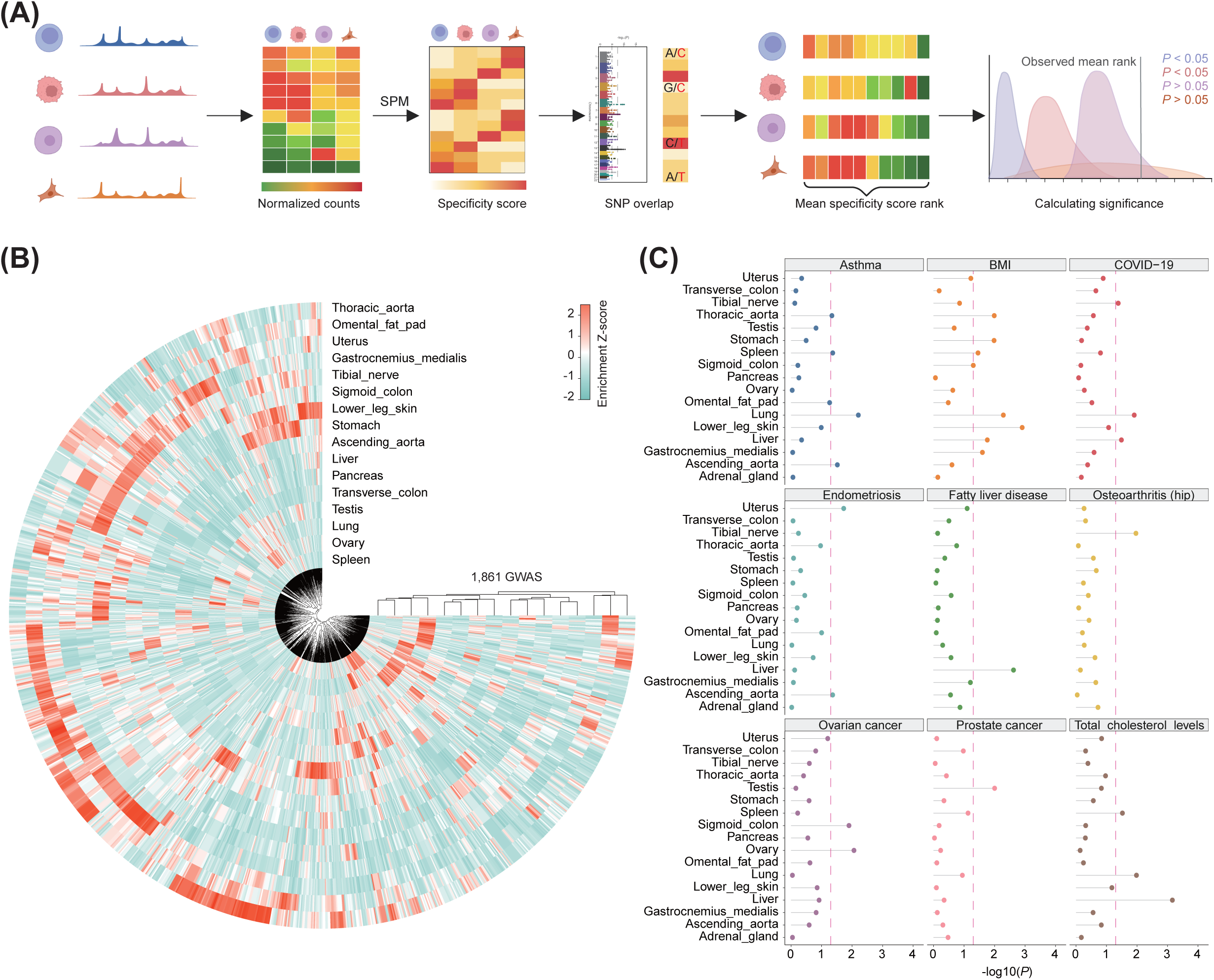
Inferring tissue types associated with complex diseases using cisDynet. (A) Schematic illustration of the principle of cisDynet enrichment for GWAS variants. (B) The circular heatmap demonstrates that 1,861 GWAS traits are significantly enriched in at least one tissue. (C) The lollipop plot showing the enrichment of nine GWAS-associated SNPs in OCRs in different tissues. The dashed line represents *P* = 0.05.

## CONCLUSION

Chromatin accessibility sequencing is a powerful technique that allows us to identify open regions within the genome, providing valuable insights into regulatory elements and enabling the construction of intricate regulatory networks. Completing these analyses often requires the use of a variety of tools, but the tools are independent of each other and need to be compatible with the running environments of the various tools, which creates a certain amount of frustration for the researcher. We have developed the cisDynet to provide a comprehensive analysis process that covers the entire process from data pre-processing to downstream advanced analysis. We provide a detailed usage guide that includes most of the usage scenarios for processing chromatin accessibility data. Notably, our cisDynet encompasses features not currently available in other toolkits, such as tissue/cell type-specific analysis, fitting of time-course data, co-accessibility analysis and integration of RNA-seq data to associate OCRs and genes. In addition to analysing chromatin accessibility data, cisDynet is also useful for analysing other epigenomic data analyses such as ChIP-seq, CUT & Tag and others. We believe that cisDynet will help researchers to accelerate the study of epigenomics.

## METHODS

### Pre-processing chromatin accessibility data

For raw FASTQ files, we first used Trimmomatic [45] to remove low-quality and potential sequencing adapters. We then used BWA MEM [46] to align the clean reads to the reference genome. We then use SAMtools [47] to convert the SAM file into a BAM file. Then the markdup function in SAMtools was used to remove the PCR duplicate fragments. Then remove the reads with low mapping quality, the default is to remove the reads with mapping quality score < 30. For BAM file with PCR duplicates removed and filtered for low mapping quality, we call it a Q30 bam file. We used MACS2 [48] software to call peak and the user can modify the necessary parameters such as effective genome size, extsize and shift as required. We finally used the bamCoverage function in deepTools [49] to generate the visual bigwig (bw) file, with default parameters smoothLenght = 20, binSize = 5. Based on the data and files obtained above, we calculated the mapping rates, PCR duplication rates, number of Q30 reads, organelle mapping rates, TSS enrichment score, FRiP, and other QC metrics, respectively. For UMI-ATAC-seq data [16], our analytical process was referred to our previously published study. We developed the data processing procedures here based on snakemake [14] and used MultiQC[50] to integrate these analyses and generate html report.

### Quality control chromatin accessibility data

For insertion of fragment distributions, we used Q30 bam files and only consider the read lengths less than 900 bp. For the calculation of TSS enrichment scores, in brief, we extracted the TSS position coordinates of all protein-coding genes, then counted the distribution of the number of reads in the upstream and downstream 3 kb of all TSSs. Finally, we used the reads in the regions of [-3000, -2500] and [2500, 3000] as the background for calculating their enrichment scores. We calculated the ratio of the number of reads falling within peaks to the total number of reads as the FRiP.

### Blacklist generation for plants

To identify blacklists on plant genomes, we followed the Greenscreen process to complete this analysis. Firstly, we extracted the BAM files of all control samples of ChIP-seq data in the ChIP-Hub database. Then peak calling was performed with MACS2 using the following parameters: --keep-dup auto -f BAM --nomodel –broad --extsize 150 --shift 75. For different species, the gsize parameter was set to Arabidopsis: 101274395, rice: 318300554, maize: 1756464757, soybean: 806805877, and tomato: 700352434. We then filtered out peaks with log10(qvalue) < 10, and then proceeded to merge all the peaks and counted the frequency of occurrence of each peak in all samples using the BEDtools merge function. Finally, we consider a peak that occurs in more than half of the input samples to be a “problematic” region.

### Differential peak analysis

Before differential analysis, we need to get a quantitative matrix first. Briefly, we first use the bed_merge function in the valr [51] package to merge the obtained peaks of different samples to get a peak superset, and then move the reads of Q30 according to the positive chain +4, negative chain -5 to get the real Tn5 cuts. Then bed_map in valr [51] package counts the number of Tn5 cuts of different samples in each peak. Normalization was then done using the normalize.quantiles function in the preprocessCore R package (https://github.com/bmbolstad/preprocessCore), divided by the length of each peak, and finally adjusted the number of cuts in each sample to one million. For the analysis of the difference peak between the two groups, we used DESeq2 [52] to identify the difference peak. For the three-group differential peak analysis, we first define a classification matrix and then assign the type of that peak according to which classification the Euclidean distance is closest to. Determination of target genes for differential peaks was done according to the TSS of the peak’s summit to the nearest gene.

### GO enrichment analysis

GO enrichment analysis is done by the enrichGO function in clusterProfiler [53]. The GO pathway enrichment difference between different clusters is done by the compareCluster function in clusterProfiler [53].

### Tissue or cell type specific peak analysis

For tissue-specific peaking analyses, we referred to previously published studies and made some additions. We first used the philentropy R package on the quantified and normalized matrix to calculate the Shannon entropy for each peak. Tissue specificity scores (TS) were then calculated according to the following formula (1). A higher tissue specificity score means a higher tissue specificity. Then the top 10% or 20% of the peaks were selected as tissue-specific peaks according to the distribution of tissue-specific scores. In addition, we calculated specificity measure (SPM) scores based on formula (2) to help us select the more specific peak in each tissue.

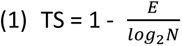

where TS represents the tissue specificity score, *E* represents Shannon entropy, and *N* represents the number of samples.

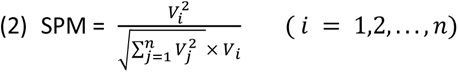

where *n* represents the number of samples and *V* represents the quantitative value of peak for the corresponding sample.

### Time-course data analysis

We begin by selecting peaks that change dynamically over time in the same way as above. If there are biological duplicates here, we do not merge the duplicates and then sort that quantitative matrix from left to right in time order. We then refer to this study to fit the time course data. We used the Local Polynomial Regression Fitting model to fit the dynamics of individual peaks at different time points, and then inserted 2,000 pseudo-data uniformly between these time points, the values of which were predicted by the model. Then we scaled the values of each row to get the *Z*-scores and visualized it with ComplexHeatmap[54]. In order to obtain the dynamics of the expression of the target genes of the peaks, we used the method mentioned below (linking peaks to genes) to get the most correlated genes for each peak. The default correlation coefficient threshold is 0.5 and the nearest one is selected as the target gene.

### TF Footprint analysis

For the TF footprint analysis, we applied the TOBIAS [21] process. Initially, we corrected Tn5 preferences by using the ATACorrect function in TOBIAS. Following that, we assessed potential TF binding sites by using the FootprintScores function to score footprints. Lastly, we identified differential footprints by using BINDetect. To compare the activity of certain TFs in specific samples, we obtained all feasible TF binding sites and extended them by 200 bp on the left and right. We then utilised the normalizeToMatrix function in the EnrichedHeatmap [55] R package to determine the distribution of corrected Tn5 cuts in that region.

### Co-accessibility analysis

After obtaining the normalized quantitative matrix, we used Cicero [56] to assess the co-accessibility between peaks. We only consider the co-accessibility of peaks on the same chromosome, and we first set the window size (default is 500 kb) to decide whether to calculate the co-accessibility scores of two peaks based on whether their summit distance is less than the window size. For peaks that fall within the same window, the co-accessibility score (from 0 to 1) between the two peaks is calculated. They are then classified as low, medium and high according to their scores. According to their original peak annotations, they were classified into promoter-promoter, enhancer-enhancer and promoter-enhancer interactions.

### Integration of RNA-seq

We first used RSEM[57] software to quantify gene expression and then used the TPM method to normalize to obtain a gene expression matrix. For the few samples where ATAC-seq and RNA-seq were integrated, we assigned regulatory regions to the nearest genes. The target genes of the differential OCRs were then assessed to see if the corresponding differential expression patterns were also present. For multiple samples, users can calculate Pearson correlation coefficients between OCRs and gene expression across samples to determine OCR and gene associations (See below for details). We then go on to construct regulatory networks based on this, depicting dynamic OCRs and gene expression changes.

### Linking peaks to genes

Peak-to-gene analysis requires entering the quantitative matrices of both peak and gene and ensuring that the column order of both is the same. We first filtered out low-expression genes, and retained genes with TPM values in all samples summed up to be greater than or equal to 0.1. For each remaining gene, we regarded the peaks within 50 kb (default, adjustable) upstream and downstream of the TSS of the gene as potential regulatory regions. Then Pearson correlation coefficients were calculated for these peaks and gene expression separately. To control for false positives, we randomly selected 10,000 peaks and calculated the correlation coefficients between peak and expression. These obtained pseudo correlation coefficients were used as background. Then we used the *Z*-test method to test the significance of each peak-to-gene and obtained the *P*-value. Finally, we then correct these *P*-values using the FDR method. We considered Pearson correlation coefficients >= 0.4 and FDR < 0.05 as significant peak-to-gene links.

### Building regulation network

The peak-to-gene analysis allows us to determine which peaks regulate the gene of interest. After we identify TF footprints in these associated peaks, we can establish TF-to-gene regulatory relationships. In this network, both positive and negative regulation levels are included.

### GWAS variant enrichment analysis

We refer to the CHEERS [42] process to complete the enrichment of GWAS variants in different sample peaks. First, we sorted the normalized quantification matrix by the sum of the quantification values for each peak, removing the bottom 10% (optional) of peaks, which we considered unreliable peaks. We then calculated the SPM matrix mentioned above, where each value represents the specificity score of that peak in the corresponding sample. For each sample, the peaks were sorted in descending order by specificity score. We determined whether the input SNP locus overlapped with the peaks. After obtaining the overlapping peaks, we calculated the mean of the rank of the peaks in each sample and then used the pnorm function in R to test for significance. Finally, we considered a *P* value < 0.05 to represent that the peaks in that sample are significantly enriched with that GWAS association to variants.

### Visualization

All visualizations are implemented using ggplot2 or its extension packages. Heatmaps were done using the ComplexHeatmap R package[54]. Genome tracks visualizations were done using the R package transPlotR (https://github.com/junjunlab/transPlotR). The clustering tree is implemented by ggtree[58]. Circular heatmap(Figure 6B) implemented using imageGP[59].

## CODE AVAILABILITY

The Snakemake-based pre-processing pipeline is available at https://github.com/tzhu-bio/cisDynet_snakemake. The codes of R package cisDynet are available at https://github.com/tzhu-bio/cisDynet. Detailed instructions for using the R package cisDynet are available at https://tzhu-bio.github.io/cisDynet_bookdown/.

## AUTHOR CONTRIBUTIONS

D.C. and T.Z. conceived and designed the project. T.Z., X.Z., Y.Y., Z.H., L.W. conducted the bioinformatics analysis. D.C. and T.Z. wrote the paper. All the authors reviewed and approved the paper.

## ACKNOWLEDGMENTS

This work is supported by grants from the National Natural Science Foundation of China (No. 32070656). The authors acknowledge the Center for Information Technology and the High-Performance Computing Center of Nanjing University for providing high performance computing (HPC) resources.

---

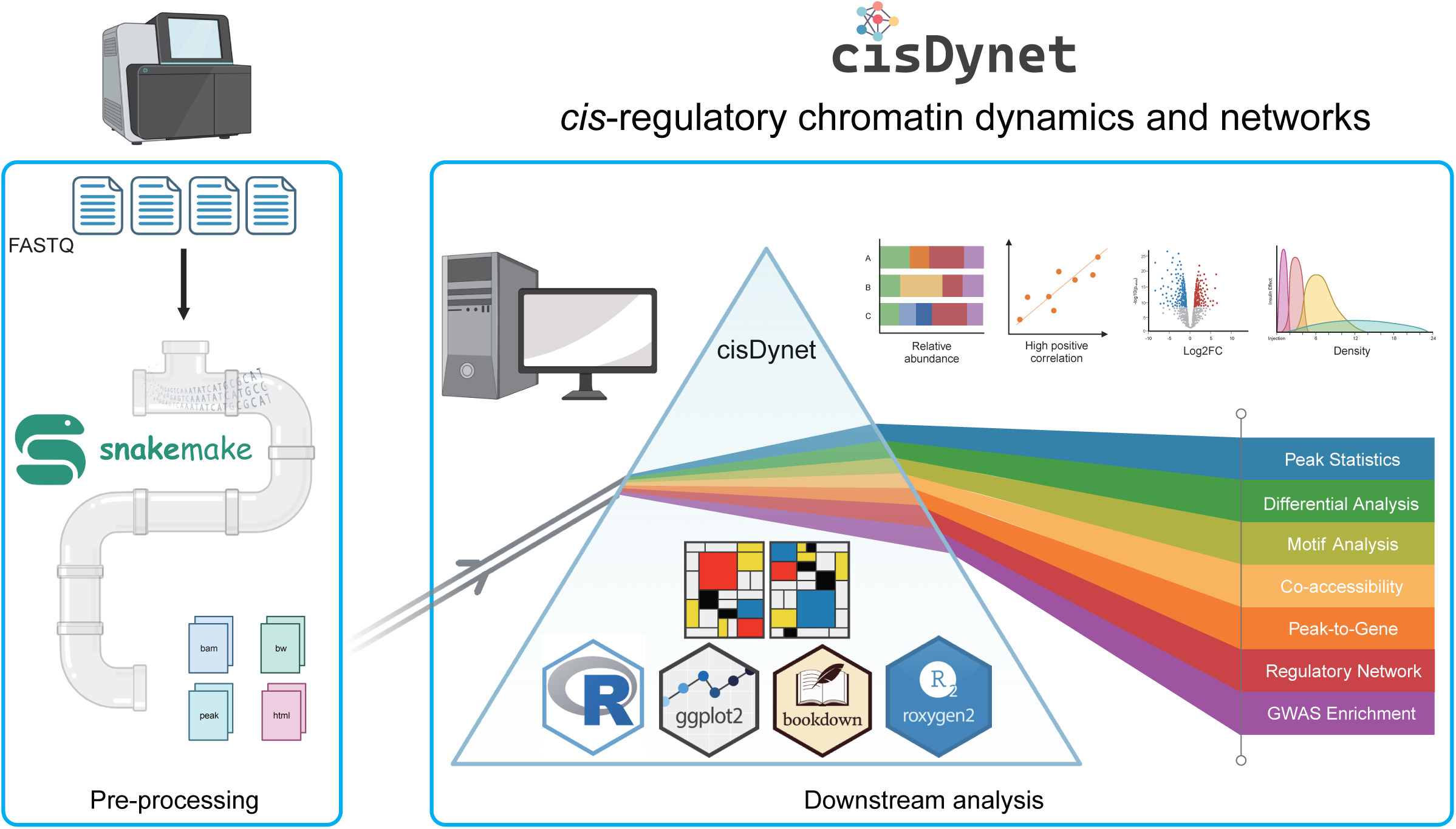

